# EasyOmics-A graphical interface for population-scale omics data association, integration and visualization

**DOI:** 10.1101/2024.02.20.581292

**Authors:** Yu Han, Qiao Du, Yifei Dai, Shaobo Gu, Wei Liu, Mingjia Zhu, Landi Feng, Jianquan Liu, Yanjun Zan

## Abstract

The rapid growth of population-scale whole genome resequencing, RNA sequencing, bisulfate sequencing, metabolomics and proteomic profiling has led quantitative genetics into a big omics data era. Performing omics data association analysis, such as genome, transcriptome, proteome and methylome wide association analysis, and integrative analysis on multiple omics dataset requires various bioinformatics tools rely on advanced programming skills and command-line environments, which are challenging for most wet-lab biologists. Here, we present EasyOmics a stand-alone R Shiny application with a user-friendly interface for wet-lab biologists to perform population-scale omics data association, integration and visualization. The toolkit incorporates multiple functions designed to meet the increasing demand for big omics data analyses, ranging from data quality control, heritability estimation, genome wide association analysis, conditional association analysis, omics quantitative trait locus mapping, omics wide association analysis, omics data integration and data visualization etc. A wide variety of publication quality graphs can be prepared in EasyOmics using R plot engine with point-and-click. EasyOmics is a platform-independent software that can be run under all operating systems with a docker container for easy installation. It is freely available to non-commercial users at docker hub https://hub.docker.com/repository/docker/yuhan2000/gwas/general.

## Introduction

With the rapid development of population-scale whole genome resequencing, RNA sequencing, bisulfate sequencing, metabolomics and proteomic profiling, quantitative genetics has entered into a big omics data era. Population-scale association analysis became more and more popular in dissecting complex traits in human, plant, animal and bacteria (Uffelmann et al., 2021). GWAS discoveries significantly improved our understanding on genetic regulation of complex developmental, disease and agronomic traits. However, the majority of GWAS signals are located in non-coding regions, complicating the fine mapping process and biological interpretation. Omics data, such as gene expression or methylation are mediators between genetic variation and phenotype. Incorporating omics data into association analyses has offered unique opportunities to interpreting the GWAS signals and revealing biological mechanisms underlying complex traits (Barbeira et al., 2021; Battle et al., 2015; Grubert et al., 2015; Hannon et al., 2015; Liu et al., 2022). Advances in association analysis have led to the development of many software tools to performing population scale omic data association analysis, such as genome wide association analysis, expression and epigenetic quantitative traits mapping, transcriptome wide association analysis and integrative analysis on multiple omics dataset to perform causal inference (Chen et al., 2019; Delaneau et al., 2017; Gusev et al., 2016; Jiang et al., 2019; Marchini et al., 2007; Zhang et al., 2019; Zhou & Stephens, 2012; Zhu et al., 2016). However, performing such analysis requires various bioinformatics tools rely on advanced programming skills and command-line environments, which are challenging for most wet-lab biologists. Therefore, the growing demand for population-scale omics data association, integration and visualization calls for a efficient graphical interface that integrates major functionalities of omics data analysis that allow simple point-and-click.

Here, we developed EasyOmics, a graphical interface for population-scale omics data association, integration and visualization. It is built on R Shiny framework and incorporates nine functions to perform data quality control, heritability estimation, genome wide association analysis, conditional association analysis, omics quantitative trait locus mapping, omics wide association analysis, omics data integration and data visualization etc. A wide variety of graphs can be prepared in EasyOmics using R plot engine with point-and-click to meet the demand to analyze and visualize growing omics data analysis. EasyOmics is a platform-independent software that can be run under all operating systems with a docker container. It is freely available to non-commercial users at docker hub. EasyOmics can significantly simplify population-scale omics data association, integration and visualization analysis process, which will be of great interests to a wide range of biologists.

## Implementation

### Data Matching

The “Data Matching” function prepares input files for subsequent analysis. First, EasyOmics extracts individual id from phenotype file and variant call format (VCF) files, excluding individuals lacking phenotype or genotype data. Then, Plink (Chen et al., 2019) was used to filter the VCF file for minor allele frequency greater than 0.03 and pairwise r^2^ smaller than 0.99 within 20kb windows.

### Phenotype Analysis

The “Phenotype Analysis” function provides insights into the input data and facilitates the quality control of outliers. For multi-trait analyses, EasyOmics calculates phenotypic variance, genetic variance, and estimating narrow-sense heritability using the AI-REML method. This method computes heritability from following 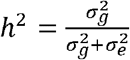 (J. Yang et al., 2010, 2011). For single trait analysis, EasyOmics presents the phenotype distribution and population structure (Chen et al., 2019) of the genotype data in a scatter plot, annotating each point with individual id.

### Genome-wide Association Analysis, Conditional Association Analysis

The “GWAS” function perform genome wide association analysis between uploaded genotype and phenotype using a linear mixed model. This approach corrects for population stratification using an identity-by-state (IBS) kinship matrix estimated with genome wide markers (J. Yang et al., 2011, 2014). In addition, it performs genomic controlled, when the inflation factor is greater than 1.1.

Moreover, conditional & joint association analyses (“COJO” function) from GCTA are integrated into EasyOmics to detect secondary association signals (J. Yang et al., 2012).

### Manhattan Plot, QQ plot and Locus zoom Plot

Manhattan plot visualize association results for all the tested SNPs. The x-axis represents the genomic position, while the y-axis displays the −Log10*(P value)*. EasyOmics offers customizable significance threshold or Bonferroni-corrected thresholds. The lead SNP will be highlighted using a red triangle with labeled id name. QQ plot present the expected distribution of test statistics for all SNPs on the x-axis against the observed values on the y-axis. Inflation factor will be calculated to control the population stratification. Locus zoom plot zooms in selected regions of interests, presenting the association results and linkage disequilibrium information. Linkage disequilibrium between selected SNPs and nearby SNPs is calculated using LDBlockShow, which also displays the results of the association analysis and gene positions using the “Locus Zoom” function (Dong et al., 2021).

### Omic Quantitative Trait Locus mapping

The “Omic QTL” function preforms omics QTL mapping using R package MatrixEQTL (Shabalin, 2012). In this analysis, omic traits, such as gene expression, epigenetic modification or metabolites profile, could be treated as molecular phenotypes. It expands our ability to perform omic data association in EasyOmics. To control for confounding with population structure, the first 20 principal components (PCs) from IBS kinship are included as covariates.

### Mendelian randomization

The “Mendelian randomization” function performs causal inference between SNPs, molecular phenotypes and traits, using genetic variants as instrumental variables. It supports analysis of two traits (e.g., flowering time as the outcome and bolting as the exposure in the “Two Traits MR” function) and accommodates gene expression or other omics data as exposure factors for rapid analysis (in the “SMR” function) (Zhu et al., 2016, 2018).

### Omic-wide Association Studies

In the “OmicWAS” function, we used a mixed linear model-based omic association (MOA) approach implemented in OSCA to test for associations between omic data and a complex trait (Zhang et al., 2019). The covariance structure of the random effect was defined as the omic-data-based relationship matrix (ORM), which captures the molecular phenotypic similarity between pairs of individuals.

### Availability

The source code and tutorial were available at GitHub repository: https://github.com/HanYu-me/EasyOmics. Its developer version is available at docker hub repository: https://hub.docker.com/repository/docker/yuhan2000/gwas/general.

## Demonstration of EasyOmics Framework and Major Functionalities

### Panel and workflow

EasyOmics is an R Shiny application with a user-friendly graphic user interface (GUI) for population-scale omics data association, integration and visualization. It implements multiple functions to perform data quality control, heritability estimation, genome wide association analysis, conditional association analysis, omics quantitative trait locus mapping, omics wide association analysis, omics data integration and data visualization etc. Each function is independent of each other, allowing users to inputting data, adjusting parameters, running analysis, and visualizing the results by simply pointing and clicking (Fig 1A). EasyOmics can significantly simplify population-scale omics data association, integration and visualization analysis, which will be of great interests to a wide range of biologists (Fig 1B).

**Fig 1.**
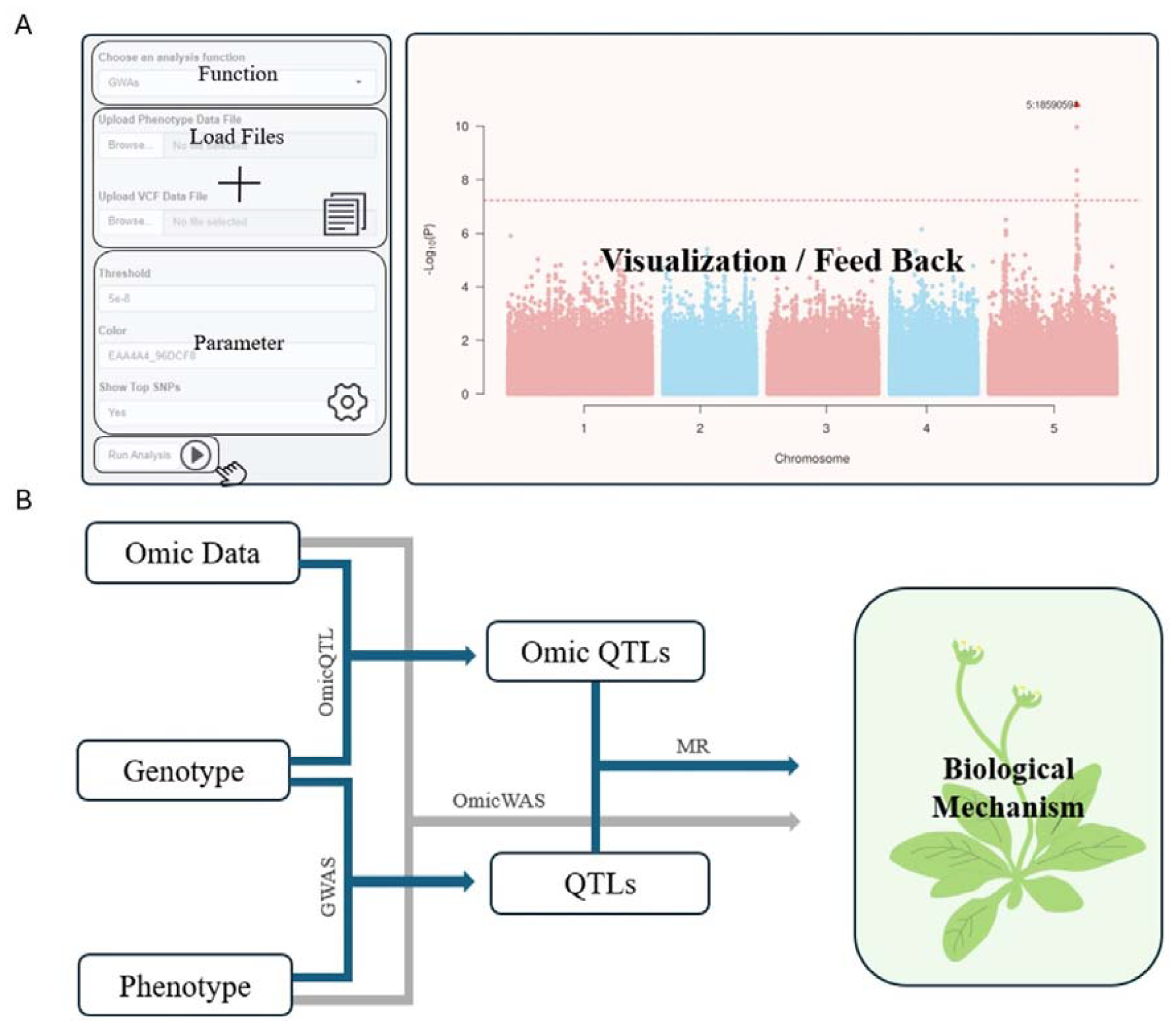
**A)** Schematic illustration of the EasyOmics application panel. The sidebar contains a selection list for analysis functions, a file upload panel, a parameter control panel, and an action button. The main panel visualize analysis output and display feedback. **B)** Schematic illustration of EasyOmics workflow. GWAs function can perform association analysis between genotype and phenotype and find QTLs with significant association. Omics QTL function treats omic data as molecular phenotype and tests the association with genotypic data. MR function integrates the QTLs and OmicQTLs to perform causal inference. OmicsWAS tests the association between phenotypic and omics data.

### Demonstration of EasyOmics

Here, we demonstrated major functionality of EasyOmics on a Windows 11 system with Intel Core i5-8265U CPU 1.60GHz and 16 Gb RAM.

We used a demo data with 10 flowering time measurements for 512 wild *Arabidopsis thaliana* accessions, collected from two common gardens (TXX or MXX located in Tübingen or Madrid) with variable drought (XHX or XLX with high or low water availability), competition (XXP or XXI with high or low plant density in pot), and temperature (FT10 or FT16 at 10°C or 16°C greenhouse) stresses (Consortium, 2016; Exposito-Alonso et al., 2019). After preprocessing the demo data with “Data Match” function, “Phenotype Analysis” function was used to summarize and visualize the data.

Flowering time displayed continuous variation from 73.67 days (FT16) to 165 days (TLP) with variable phenotypic variation across environment. There was moderate variation (0.34-0.90; Fig 2A) in the estimated phenotype narrow-sense heritability from one environment to another. Consistent with this, the amount of additive genetic variation (V_a_) estimated as V_p_ times *h*^*2*^ changed considerably, highlighting existence of phenotypic plasticity or genotype by environment interaction (Ji et al., 2023) (Fig 2A). Taking FT16 as an example, EasyOmics displayed phenotype distribution and population structure of each accession with labelled ID, which helps the user to quality control the input data (Fig 2C, D).

**Fig 2.**
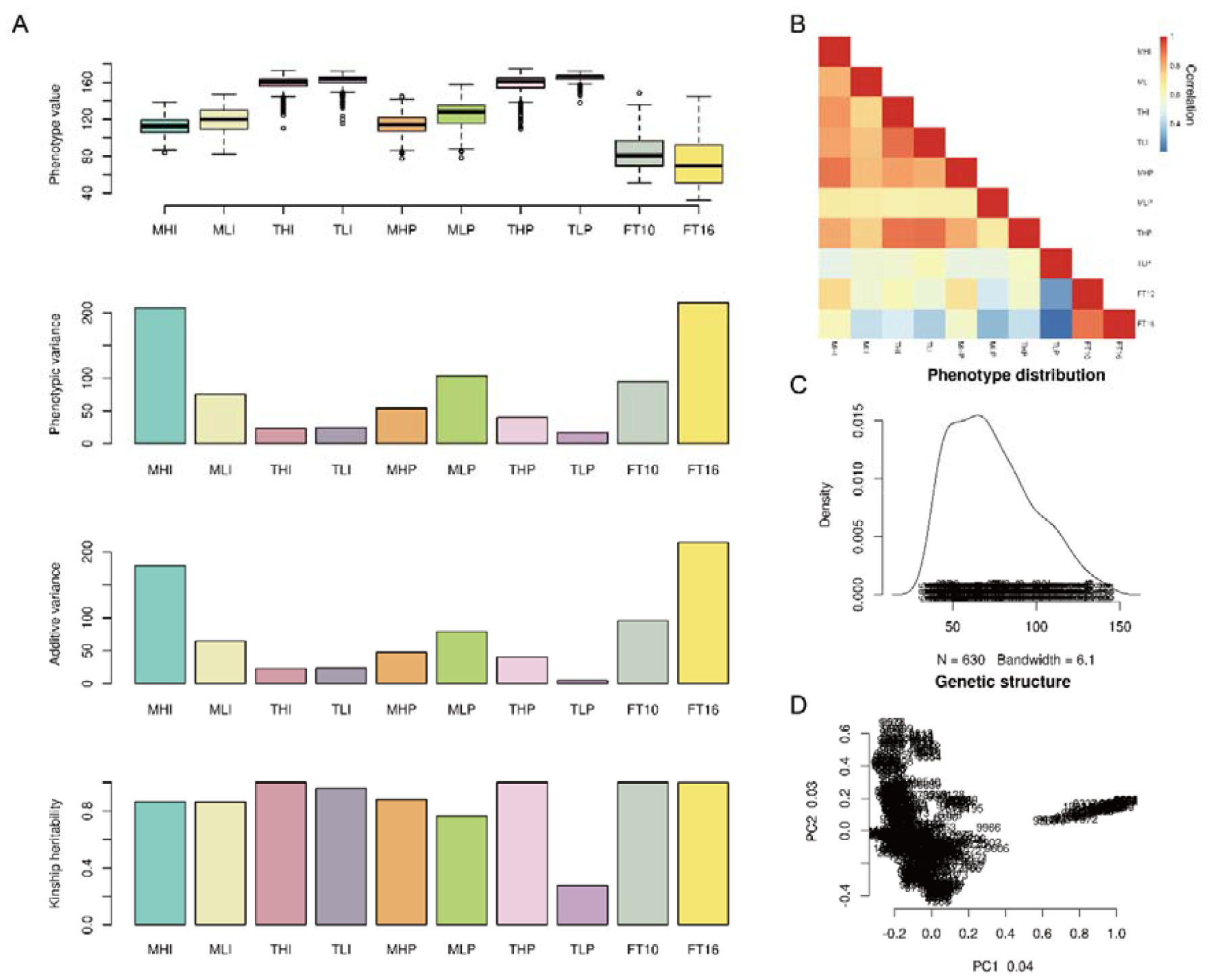
Output from the “Phenotype Analysis” function. **A)** Boxplot illustrating multi-site phenotype distribution, the phenotypic variance and additive variance, as well as the heritability estimation. **B)** Pairwise Pearson correlation among input phenotype. **C)** Density plot of selected phenotype, accessions were annotated by their accession number at the bottom of the figure. **D)** Population structure visualized as first two PCs calculated by the genotype data for all accessions.

### Genome wide association scan, conditional scan, manhattan plot, QQ plot and Locus Zoom plot

Here, we performed a genome-wide association analysis utilizing the “GWAs” function implemented with a mixed linear model to detect genetic variations associated with FT16. It automatically outputs a manhattan plot and a QQ plot in the working panel. In addition, this function priorities association signal by filtering the significant SNPs based on statistical significance, physical distance, and Linkage Disequilibrium (LD). In this case, this analysis prioritized QTL 5:18590591 (P = 1.58 x 10^−11^) as the lead SNP to represent this association peak (Fig 3A). The inflation factor was estimated to be 1.02, indicating confounding with population stratification was fully adjusted (Fig 3B). Otherwise, genomic controlled will be performed to further adjust the confounding with population stratification.

**Fig 3.**
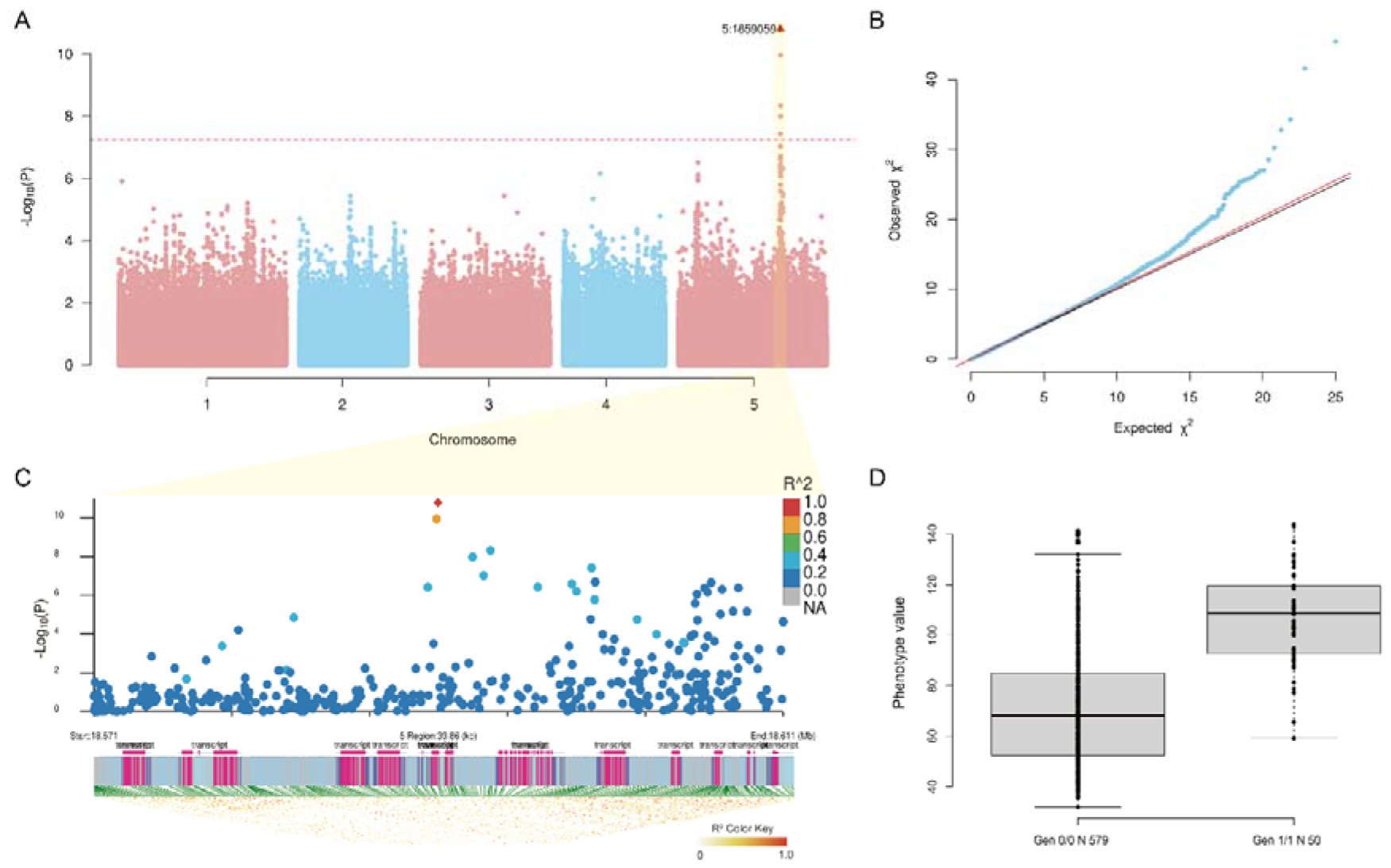
Output from GWAS related functions. **A)** The Manhattan plot. The horizontal dashed red line is the genome-wide significance threshold after Bonferroni correction, and lead SNP was highlighted using a red triangle with labeled id. **B)** The QQ plot. **C)** Regional association landscape and LD heatmap around a 20 kb region of the lead SNP. The top panel displays a regional manhattan plot around the lead SNP, where the color represents the LD gradient with the lead SNP. The middle panel displays the gene annotation. The bottom panel displays the LD heatmap, and the color of the grid represents the degree of LD measured by R2. **D)** The boxplot indicates the genotype and phenotype map at this locus.

The “Locus Zoom” function presents regional association landscape and LD heatmap underlying the lead SNP within a 20 kb genomic region (Fig 3C). In the demo data, eleven genes were located in this region, including *DOG1* characterized with flowering-related function (Consortium, 2016; Huo et al., 2016; Martínez-Berdeja et al., 2020; Taylor et al., 2017). At this QTL, accessions with the alternative genotype flowered significantly later than those carrying the reference genotype (Fig 3D). Additionally, Conditional association analysis was performed in “COJO” function, and failed to reveal any additional loci directly associated with FT10.

### OmicQTL detect genome wide cis- and trans-QTL profile for gene expression regulation

The “OmicQTL” function perform association analysis between genetic variants and omic data, such as transcriptome, methylome and proteome. We demonstrated the functionality of this function using a public transcriptome dataset with 728 *A*.*thaliana* accessions (Kawakatsu et al., 2016). After processing data using “Data Matching” and”OmicQTL” function, 630 accessions and 22,119 genes with detectable RPKM values in more than 50 % accessions were retained.

In total, 190,950 SNPs were detected with significant associations to the variation of 9,872 genes (P < 1×10^−8^). They were further merged to 14,872 eQTL by only including lead SNPs. The majority of genes were detected with cis-eQTL, while two trans-eQTL hotspots were found in chromosomes 4 and 5 (Fig 4). One of the hotspot located on chromosome 5:22,583,967 bp to 22,605,170 bp is simultaneously associated with the expression of 156 genes, including 33 flowering time genes (Zan & Carlborg, 2018). Within 20 kb, we found a well-known candidate gene, AT5G55835. It is a microRNA targeting several SPL family members, which plays a role in regulating various developmental processes, including the transition from vegetative to reproductive growth and abiotic stress (Wang et al., 2009; L. Yang et al., 2011).

**Fig 4.**
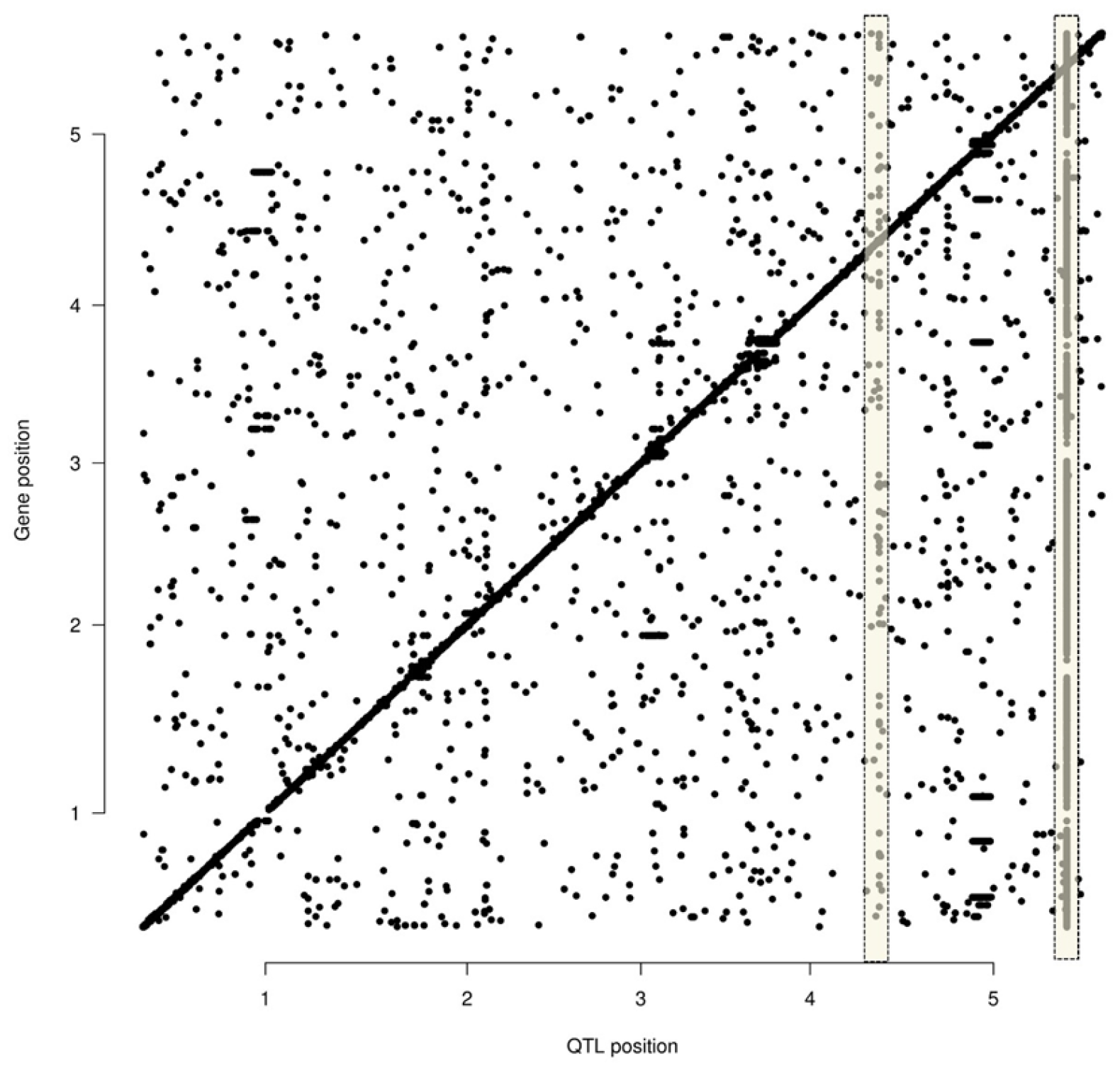
Genome wide cis- and trans-eQTL landscape, where X-axis is the location of eQTL, and y-axis is the position of associated genes. Light yellow bars indicated trans-eQTL hotspots.

### Mendelian randomization bridges the gap between SNP and trait association

Mendelian randomization uses genetic variant as an instrumental variable to unravel causal relationship between complex or molecular traits such as gene expression and other complex traits. Here, we treat gene expression as an exposure variable and flowering time as an outcome variable to identify the causative effect of gene expression on phenotype using the “SMR” function. Please note that the population size in our demo data is limited and most of the QTL detected in flowering time GWA scan did not show significant associations after Bonferroni correction. For illustration purpose, we reduced the significant threshold to 1×10^−4^ to demonstration the functionality. There were two genes (AT5G45730 AT1G66100) showed positive signals (Fig 5B, C). The scatter plot displays the effect sizes of eQTL on gene expression and flowering time (Fig 5B, D).

**Fig 5.**
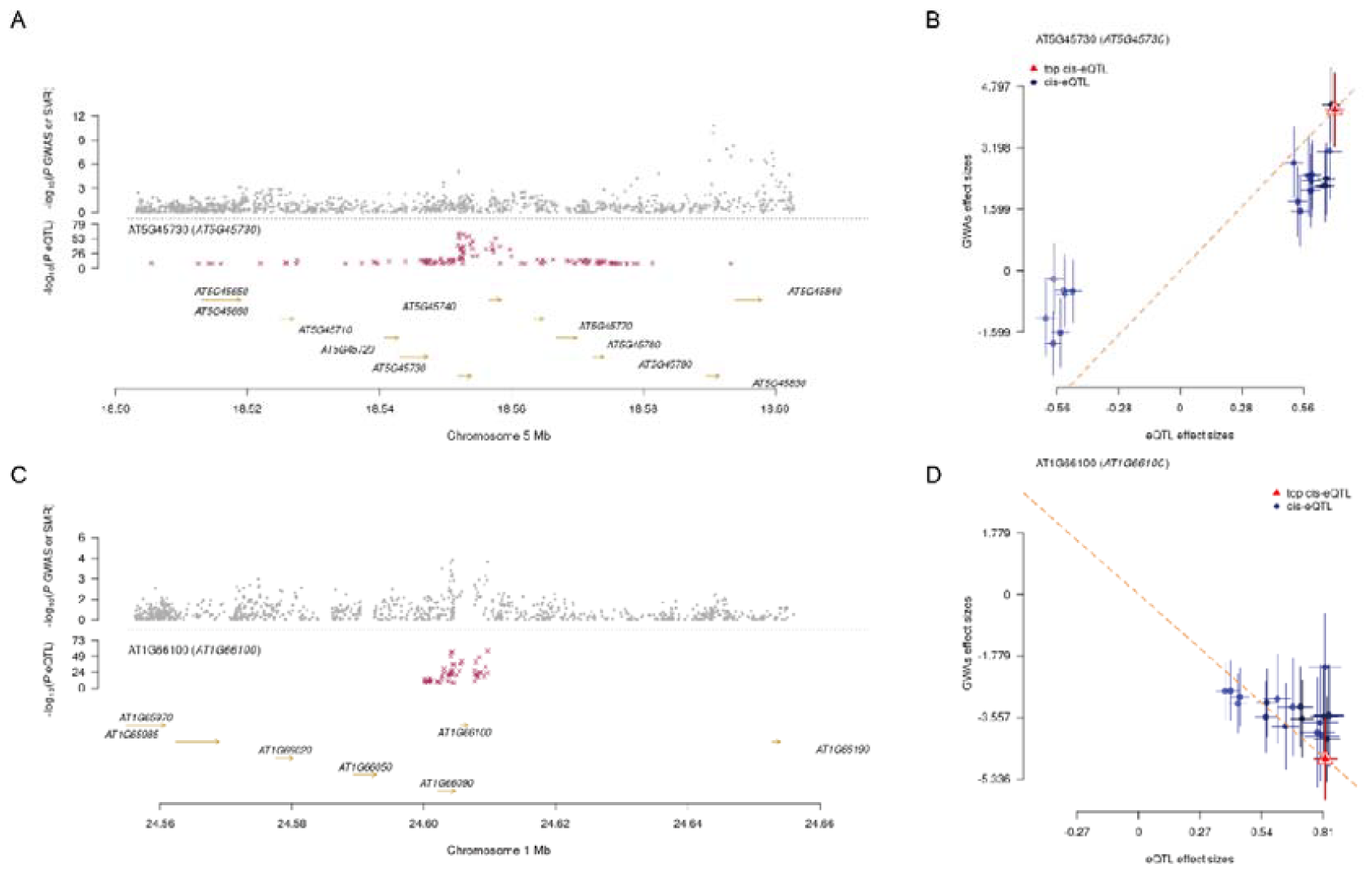
Output from Mendelian randomization analysis. **A)** and **C)** Prioritizing genes at a GWAS locus using MR analysis. Shown are results for AT5G45730 and AT1G66100 whose expression was associated with flowering time. Top panel is the P values for SNPs from GWAs. Middle panel is the P values for SNPs from omicQTL. Bottom panel is the gene position in this region. **B)** and **D)** Scatter plot of SNP effect on gene expression (AT5G45730 AT1G66100) and flowering time. The orange dashed lines represent the estimated gene effect on phenotype. The top cis-eQTL was highlighted in red. Horizontal and vertical dotted black lines indicate 95% confidence intervals for the exposure and outcome associations, respectively.

### Omic wide association analysis

We performed a transcriptome wide association anayisis (TWAS) using the “OmicWAS” function to test for associations between genetically regulated gene expression and complex diseases or traits. A total of seven genes were identified with significant association with flowering time at FT16 (Fig 6). *SOC1*(AT2G45660) acts downstream of FT, regulating flowering time and phase transitions (Lee & Lee, 2010; Quiroz et al., 2021; Taylor et al., 2019). The analysis runs at the gene level facilitates users to quickly pinpoint candidate genes.

**Fig 6.**
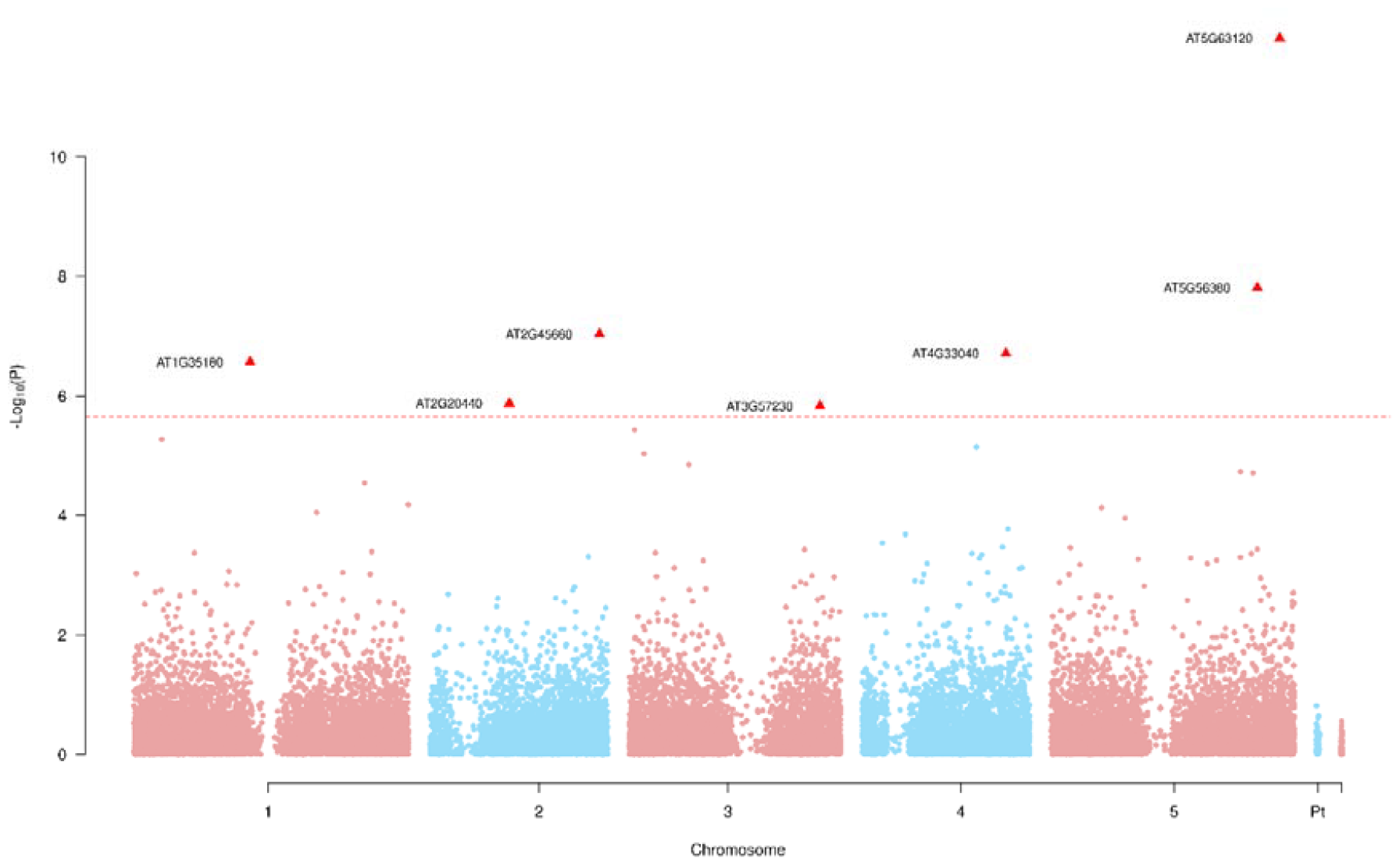
Output from OmicWAS function. The horizontal red dashed line is the Bonferroni corrected genome-wide significant threshold. Each dot represents a single gene, and significant associated genes were highlighted as a red triangle labeled with their ID.

In summary, we present a toolkit incorporates multiple functions designed to meet the increasing demand for big omics data analyses, ranging from data quality control, heritability estimation, genome wide association analysis, conditional association analysis, omics quantitative trait locus mapping, omics wide association analysis, omics data integration and data visualization etc. A wide variety of publication quality graphs can be prepared in EasyOmics using R plot engine with point-and-click. All the analysis could be completed on a laptop within reasonable time. We think the functionality, flexibility and efficiency will be of great interests to a wide range of biologists.

